# Energy-Based Systematic Modeling of Adaptive Immune Repertoire Behavior: Study on Cell Proliferation & Somatic Hypermutation process

**DOI:** 10.1101/2024.05.13.593908

**Authors:** Yexing Chen, Haiwen Ni, Jin Ma, Yongjie Li, Chen Huang, Sixian Yang, Xiangfei Xie, Haitao Lv, Min Li, Peng Cao

## Abstract

Monitoring and describing the adaptive immune repertoire(IR) for disease detection and diagnosis is essential in healthcare research. Many phenomenon have been observed to generalize the static property of IR, but the mathematical description of the formation and dynamic changes of IR still lacks research. Here, we present a mathematical and physical model to interpret the cell proliferation and somatic hypermutation(SHM) process in IR, difficulties to generate different clones in IR are computed, and IR distance is calculated as the minimum effort required to transform one IR distribution to another. IR distribution and distance can be detected from samples containing 10**^4^** cells. The validity of our method is confirmed by the unsupervised clustering of data from mice spleen and clinical PBMC samples including various immune stages and diseases. Our work dynamically characterize and quantify IR process, enabling a macroscopic immunoevaluation by sensitive immune fluctuation detection from minute samples.

## 1 Main

The adaptive immune system mounts a specific response to antigens including pathogens such as bacteria and viruses, as well as in autoimmune diseases and cancer. Adaptive immunity is triggered when a threshold dose of antigen are generated(Miller, 2000), this process may already occur before the manifestation of any clinical symptoms. Adaptive immunity exhibits distinct response characteristics under the stimulation of different antigens(Mhanna et al, 2024), so by detecting the activation and characteristic of adaptive immunity, early disease diagnosis and etiological screening may be achieved.

Antigen recognition in adaptive immune system is achieved through B cells and T cells by their respective receptors, the B cell receptor(BCR) and T cell receptor(TCR). These receptors exhibit a high degree of diversity to ensure compatibility with the diverse range of antigens(Pai and Satpathy, 2021; Miho et al, 2019). Immune repertoire refers to the concept of the functional assemblies of BCR and TCR receptor, which shifts due to the selection of high-affinity cells during the adaptive immune process. Central challenges in quantitative repertoire analysis include the dynamic description of repertoire generation(Altan-Bonnet et al, 2020), summarizing the diversity of the entire repertoire(Roswell et al, 2021), the computation of the repertoire shift(Brown et al, 2000), as well as the correlation between disease and repertoire behavior(Finotello et al, 2019). Several approaches have been developed to depict the repertoire from high throughput sequencing data, such as the diversity measurements(Roswell et al, 2021), similarity aggregating methods(Collins and Jackson, 2017), and Artificial Intelligence(Sidhom et al, 2021), some methods describe the origination of diversity during cell differentiation, while some other indicators tend to describe a transient state of repertoire. However, indicators such as Shannon Index requires data normalization due to its extensive properties related to sample size, which can introduce unnecessary systematics errors(Extended Data Fig. 1)(Hsieh et al, 2016). Moreover, these existing indicators are unable to demonstrate the direction of repertoire development. Thus, quantitative methods are still insufficient in immune repertoire research.

As the complex repertoire evolves huge diversity, it is difficult to quantify the diversity or characterize the overall repertoire status based on statistics. A similar situation exists in the study of gases in physics, where gases have a variety of velocities and directions of movement, making it difficult to distinguish between two samples by sampling gas molecules and statistically analyzing a specific characteristic. Although it is not possible to find a specific characteristic, gases exhibit stable macroscopic properties such as density, temperature, and pressure on a large scale, which are collective behaviors exhibited by a large number of molecules. Therefore, similarly, we do not study the specific characteristics of BCR and TCR gene diversity, but rather the collective behavior of a large number of cells, in order to uncover the underlying mathematical and physical laws.

So we first constructed a coordinate space to present the status of repertoire based on the results in HTS sequencing, then the difficulty of each point in the space for the immune system to occupy was simulated and calculated, finally we quantified and calculated the repertoire shift as the the minimum effort required to transform a repertoire distribution to another. After blind verification with clinical and mouse data, we believe that this method can indeed describe the macroscopic state of the entire immune system and reflect its diverse activation patterns in response to different diseases.

### 1.1 Quantitative visualization of the repertoire distribution

We first built a 3D structural demonstration of repertoire, with mutation(X axis), clone kind(Y axis) and clone proliferation(Z axis) as base substrates(Extended Data Fig. 2), where clones were defined as cells with the same BCR heavy chain or TCR *β* chain nucleotide sequence. The scale of Y and Z axis are normalized to 1. Each cell in the repertoire get its unique point occupation in the space and the cell distribution can be intuitively embodied. Previous discoveries about the repertoire can be systematically reflected in this space: The variations in the average and median mutation number within a repertoire offer a vivid reflection of the processes of SHM and antibody affinity maturation(Mesin et al, 2016), mutations that occur at a frequency exceeding the random mutational background are designated as hotspot motifs(Vieira et al, 2018), the observation that the clone exhibiting the highest frequency does not necessarily harbor the greatest number of mutations is indicative of an epitope spreading(Vanderlugt and Miller, 2002). Occupancy of various points within the 3D space engenders a spectrum of distinctive immune signatures. Therefore, we opted for this form of representation to articulate the immune repertoire.

The distinctive signatures of the immune repertoire associated with each pathology can be delineated and, following rigorous summarization and validation, can generate a multitude of bioinformatics indices. However, these indicators vary across different diseases and represent empirical findings, and cannot give quantified results. To investigate the ubiquitous principles governing immune system activation, we chose to systematically study the difference between two different repertoire distributions rather than summarizing empirical outcomes. The key question of the study is to determine the quantitative measure of effort necessitated for the immune system to assume presence at specific points within the 3D space, and how to quantify the differences between two 3D distributions.

### 1.2 Calculating the difficulty to generate a repertoire

To calculate the weight of each point in the space, we decided to employ a Markov process and conduct computer simulations to examine the immune activation mechanism. We began by transforming the uneven cell proliferation process into a preferential selection model, highly proliferative cell clones have higher probabilities to add new cell members(Fig. 1A). The emergence of preference is based on the different amplification rates caused by BCR or TCR binding capacity which is determined by VDJ recombination(Teng and Schatz, 2015) and SHM. Six parameters(four for TCR) are set in the model to describe the process(Extended Data Fig. 3), an immune repertoire distribution is uniquely determined by these parameters. (i) the distribution of initial activating signal stimulated by antigens. (ii) the initial number of B or T cells involved in a immune response, (iii) the effective or non-lethal SHM rate(Kleinstein et al, 2003) in BCR which change stimulating signal randomly, (iv) the amplitude of change in stimulating signal caused by SHM, (v) the positive selection and the formulation to calculate the proliferation speed from stimulating signal. (vi) the overall increase in the number of cells and clones through cell division and mutation.

**Fig. 1.**
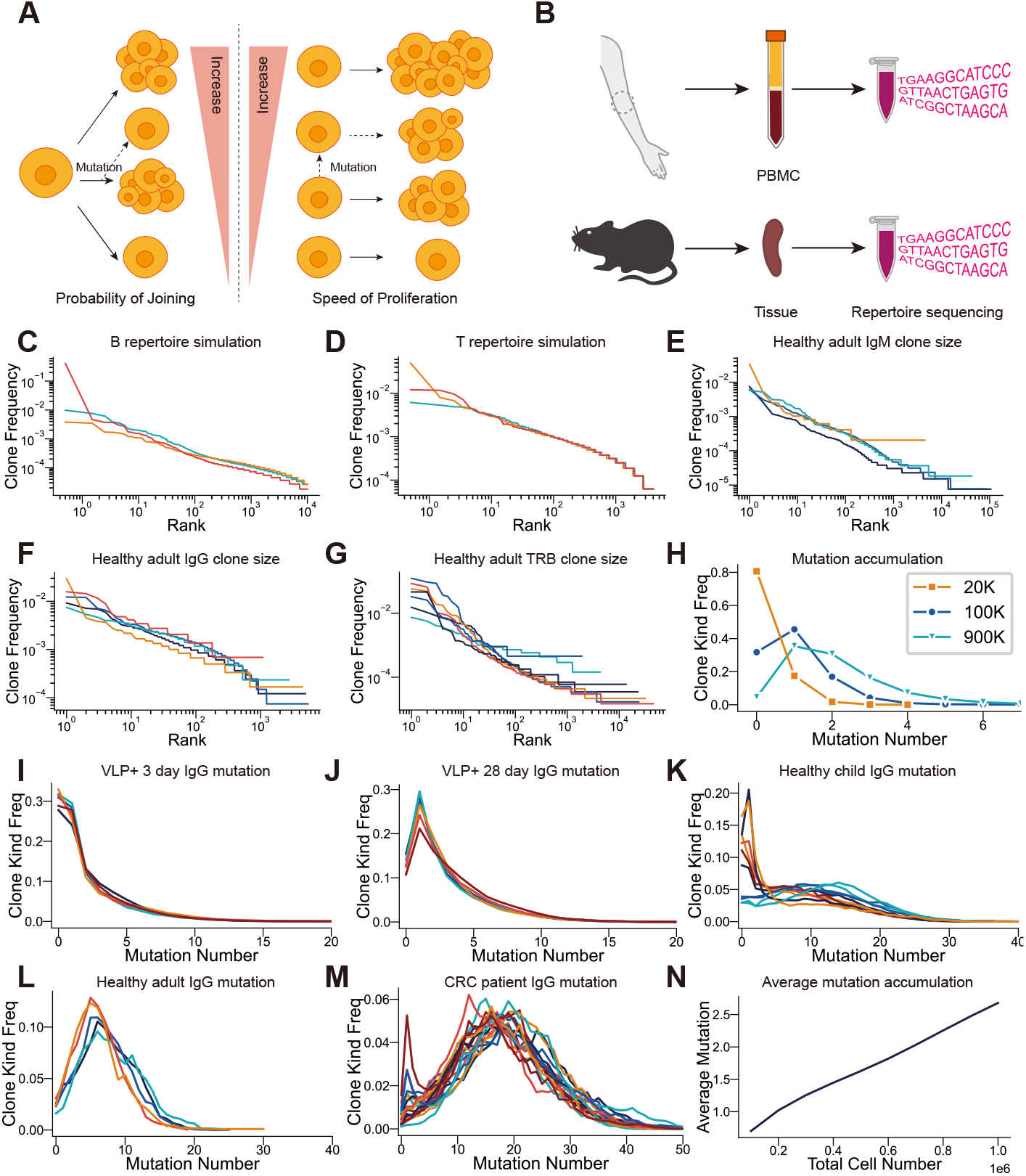
The preferential attachment model simulate the rank-frequency and mutation distribution of repertoire. Each line represents an individual or an simulation instance. (**A**) Clones with high proliferation speed equals to have high probability to attract new cells to attach to them, New clones are generated when mutation occurs. (**B**) Samples are collected from human peripheral blood and mouse spleen, bulk HTS sequencing are used to get repertoire data directly from total RNA without FACS sorting or single cell techniques. (**C, D**) BCR&TCR repertoire generated from the model show the scale free characteristics. (**E, F, G**) the scale free characteristics of IgM, IgG, and TRB repertoire data from healthy adults. (**H**) The simulated distribution of mutationclone shifts with different repertoire size, the immune process involving more proliferation has a higher average mutation distribution. (**I, J**) C57BL/6 mice are immunized with VLP, the mutationclone curve in early(3 days) and late(28 days) stages shows similar distribution with the simulated repertoire. (**K, L, M**) The mutation-clone curve in clinical samples. The unexperienced(healthy child), experienced(healthy adult) and well activated(cancer patients) immune system also show the similar unimodal and shift pattern as in the simulated repertoire. (**N**) In the simulated repertoire, mutation accumulates linearly with the proliferation process.

The progression of cell proliferation and mutation during adaptive immunity is simulated by repeating the attachment process, a starting repertoire distribution is first initialized then the amplification, SHM and selection process is duplicated cell by cell to mimic the progression of the immune response, the system’s attributes are extracted and analyzed throughout the process.

Real world data were collected to help validate the model(Fig. 1B). Spleen data were collected from C57BL/6 mice immunized with Candida Albicans Water Soluble fraction(CAWS) or Virus-like particle(VLP)(Kozlovska et al, 1993). Meanwhile, clinical data from peripheral blood were obtained consisting of healthy controls, individuals with infections, patients diagnosed with Kawasaki Disease, and patients with colorectal cancer(CRC). The samples were selected from a natural population cohort, the healthy group did not exclude participants with recent infection or vaccination history, and the disease group did not screen for chronic diseases, complications, etc. BCR heavy chain of IgA, IgM, IgG B cells and TCR*β* gene from T cells were sequenced.

The rank-frequency curve of the simulated repertoire exhibits a fat-tail power law distribution(Desponds et al, 2016)(Fig. 1, C-D), which is consistent with the B-A preferential model that produces a scale-free network(Barabäsi and Albert, 1999), this power law distribution is also observed in real data(Fig. 1, E-G) and previous research results(Mora et al, 2010). The scale-free pattern can be considered as the dimension reduction of the 3D structure on Y-Z plane after sorting(Extended Data Fig. 4). The distribution of clones with varying mutation numbers, which is also the projection of the 3D structure on the X-Y plane, distributes an unimodal curve(Extended Data Fig. 4), and both the average and median number of mutations per clone increase with the total number of cells(Fig. 1H), these trends are observed in different immune stages(Fig. 1, I-J) and health conditions(Fig. 1, K-M, Extended Data Fig. 5A) with significant consistency. The correlation between total cell number *N*_total_ and average mutation number per cell *N*_Mut_ in the model is almost linear(Fig. 1N).

The consistency of results strongly implies a functional relationship between number of mutations and clone kind, suggesting that the analysis of the immune repertoire should not only be sequence-based but may also need consideration at the clonal aspects, where the clone size could be considered as a parameter of the clone in calculating the mutation rate, proportion, and biological function weight.

From the perspective of mechanics, if we consider the total cell number after proliferation *N*_total_ in immune process as an indicator of force exerted by the immune stimulation, and the average mutation number per cell *N*_Mut_ as a displacement of the BCR gene and whole immune system, then the effort *E*_SHM_ required to generate a cell with mutation number *N*_Mut_ is approximately proportional to *N* ^2^, similar to the relationship between tension and deformation of a spring according to Hooke’s law. TCR as well as IgM antibodies in the early stages of the immune response do not occur the mutation process, so we expand the scope of application of the model to cells without mutation by adding 0.1 in *E*_SHM_ as ground energy to avoid the occurrence of zero values as in Eq. 1, ground energy pertains to the essential effort required for the maintenance of normal cellular life processes and is preliminarily and arbitrarily set to 0.1.

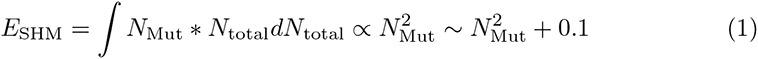

The X axis in the 3D space defines the weight of the clone, points with the same x-coordinate possess identical weight levels. The spatial distributions exhibits the direction and extent of how immune system react to different immunogenic stimuli, and can serve as an indicator for disease diagnosis.

### 1.3 Quantification of repertoire distance

After defining the weight of different clones in the repertoire, the repertoire shift now can be computationally quantified as the minimum effort required to transform a 3D repertoire distribution to another. Theoretically, the difference between two 3D distributions can be calculated by the 3D Wasserstein Distance. Due to the difficulty and complexity of computation process, in actual computation, we reduced the 3D distributions to a 2-dimensional Rank-Frequency curve then calculated the 2D Wasserstein distance(Panaretos and Zemel, 2018) by Scipy package in Python. clone weights are set as *N* ^2^ + 0.1, as calculated in Eq. 1.

The diverse functions of antibodies are implemented through distinct antibody subtypes, which can be regarded as different facets in the immune response. Therefore, sequencing data of each type including IgA, IgM, IgG antibodies and TCR*β* will generate a 3D space and a distance, total distance between two individuals are the vector sum of each distance in TCR and all antibody subtypes. Because of the class switch process, these vectors are not orthogonal to each other, at the same time, data of same subtypes are difficult to measure in clinical samples like IgE, so the distance between all accessable gene types are calculated and preliminarily use Chebyshev distance(Cantrell, 2000) to integrate them and evaluate the possible distance between two individuals.

## 2 Results

### 2.1 Unsupervised clustering of mouse and clinical data

A distance matrix was built by measuring the total distance between pairwise combinations of all biosamples, then the matrix was transformed to visualized cluster results by t-SNE. The method is unsupervised, clinical information are not used in clustering and are treated as a blind factor, the visualized clusters were labeled in the last step for correctness validation based on the clinical grouping(Fig. 2A, Extended Data Fig. 6). These samples were naturally divided into several clusters, the clusters was consistent with their clinical grouping and samples within the group exhibited strong consistency. These findings confirmed that the activation of the immune system displays distinctive patterns in response to different diseases.

**Fig. 2.**
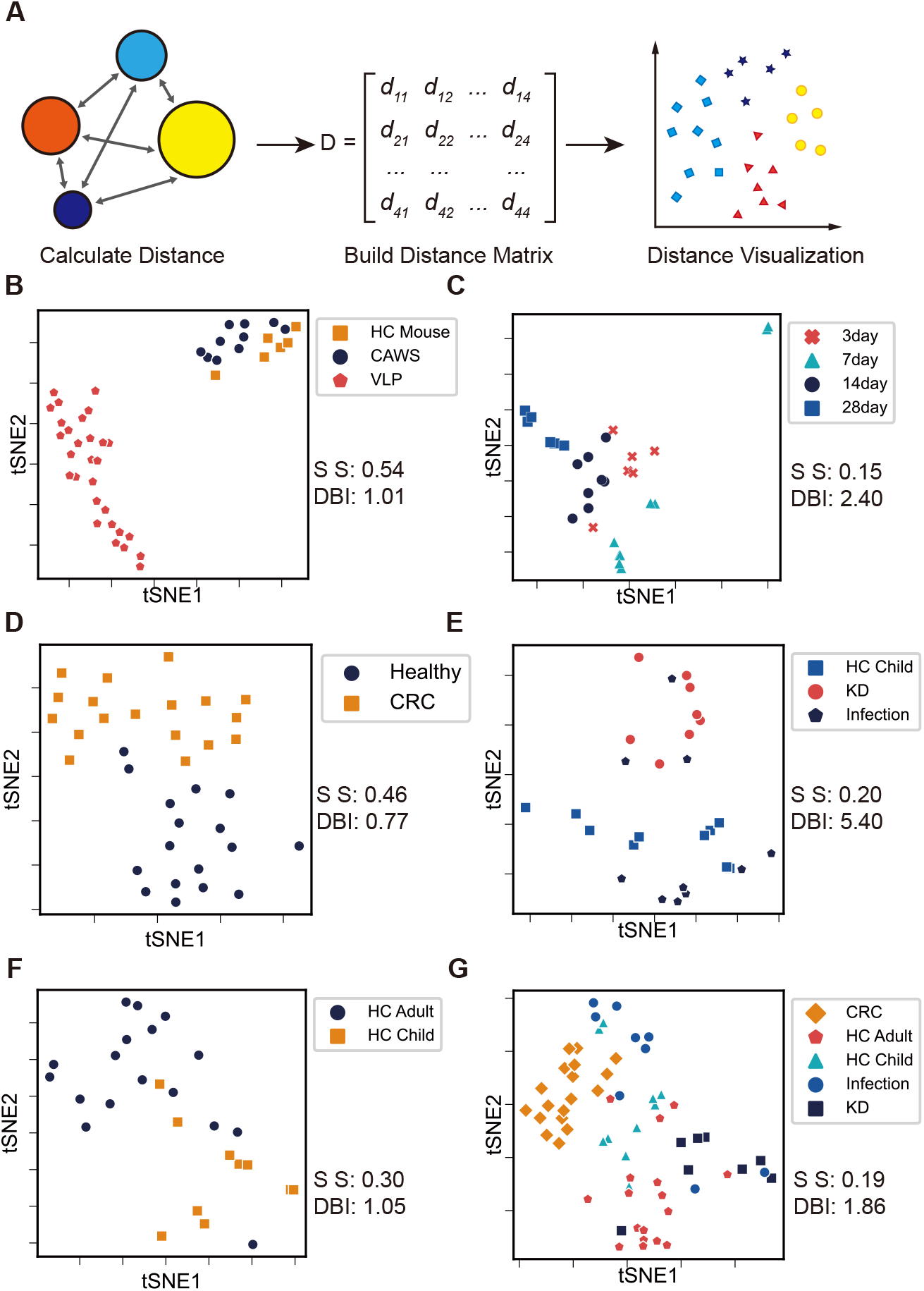
Repertoire shift calculation and sample clustering based on repertoire distance. Each point represents an individual. (**A**) Sample distances are calculated as minimum efforts required to transform one repertoire distribution to another between any two samples, distance matrix are calculated and then blindly clustered and visualized by t-SNE. (**B**) Distinct clusters are shown between C57BL/6 mice samples immunized with CAWS and VLP. (**C**) The clustering method is capable of distinguishing different immune stages in mouse VLP immunization progress. (**D**) The primary diagnosed CRC shows an unique cluster result. indicating a potential detection method for cancer early diagnosis. (**E**) The different clustering of KD and infection may help solve the dilemma of misdiagnosis and stop disease progression in early stages. (**F**) The differences in healthy child and healthy adult clustering shows that the method is also capable of depicting the dynamic change in immune system development without antigen stimulation. (**G**) Clustering of clinical samples with all healthy conditions included in the study, each condition has unique immune pattern and clusters separately.

Significant difference can be observed in mice with different antigen immunization such as CAWS and VLP(Fig. 2B), indicating that different types of antigens can trigger different immune stimulation and activation model, which lead to different immune response outcomes. Mice after VLP immunization for 3, 7, 14 and 28 days have distinct adaptive immune repertoire clusters(Fig. 2C), which proves the precision of the model in dynamic immune reaction detection during the whole immune process. With enough time point collected, it is feasible to draw the trajectory of repertoire change in the whole course of disease. The potential secondary complications and disease progression can then be detected and prevented by the deviation of the immune repertoire characteristic. These results can serve as a reference for immunological mechanism research and an identifier for the cause of disease. The VLPs capsid protein is a vector for Virus-like particle vaccines(Mohsen and Bachmann, 2022), so this method may give evaluations to vaccine effectiveness by comparing immune characteristics triggered by vaccines or real infections.

Clinical evidence also proved the effectiveness of this method. Different clusters representing different immune responces can be observed among primary diagnosed colorectal cancer and healthy controls(Fig. 2D, Extended Data Fig. 5). The two clusters showed significant differences, so we believe this can be used as an early detection indicator for colorectal cancer.

Misdiagnosis may be alleviated for some diseases in clinical practice, such as mild cases of Kawasaki disease(KD) that are easily misdiagnosed as fever caused by infection(Fig. 2E), the clustering result showed the distinct differences between two diseases, three individuals diagnosed with infectious fever, but exhibiting similar immune response characteristics as KD patients, may illustrate the existence of potential misdiagnosis in clinical practice. So the method can be used to differentiate diseases caused by different etiologies with similar symptoms, and may help aiding clinicians in the rapid identification of the cause of a disease.

The immune development over ontogenesis was observed by distinct clusters of healthy child and adult groups(Fig. 2F), thus the effect to immune system by age or other factors can be observed and researched, subhealthy status may then be distinguished and quantified by evaluating the background immune activation.

The model not only enables comparisons between pairs of different health conditions but also supports the simultaneous differentiation among a large number of various health status samples. The different clusters of all 5 health conditions in the study separated significantly to each other(Fig. 2G), so it is feasible to draw a map containing many kind of diseases and benefits clinical diagnosis unless diseases triggers similar immune activation. Since the model is constructed based on the process analysis of the normal immune system without using any specific traits from particular disease or intervention, this approach may be extensively generalizable to diagnostic and therapeutic methods across a diverse array of pathologies. Unknown samples are classified by directly comparing their calculated distances to known clusters and assigning them to the most similar cluster. Therefore, as the database of known information becomes more comprehensive, the classification of unknown samples also becomes increasingly precise.

Conventional classification models may not suitable to be applied to this method, as conventional models are only capable of identifying disease types included in the training set and lack the ability to generalize or give quantified results. Our method was unsupervised and no assumptions or statistics were introduced, so the evaluation criteria for the method should be accuracy parameters like the Silhouette score(SS)(Rousseeuw, 1987) and Davies-bouldin index(DBI)(Davies and Bouldin, 1979), rather than the false positive rate, similar to evaluating a microscope which is basing on fundamental optical principles.

### 2.2 Subgrouping within the healthy group

The healthy group did not exclude participants with a history of recent infection or vaccination(Post-IV), whose immune status may be influenced by exogenous antigens or medications. the naive healthy group consists of 5 individuals, while the Post-IV group includes 12 individuals. Significant differences in the relationship between mutation and clone frequency were observed(Fig. 1L, Extended Data Fig. 5A), and two distinct clusters could be identified in the unsupervised clustering(Extended Data fig. 5B). However, the information on the infection history of the healthy children group was not collected, the mutation distribution and clustering result within the group exhibits variability(Fig. 1K, Fig. 2 E-F, Extended Data fig. 5C). Aftering trimming healthy data, the differences between healthy and cancer have become more pronounced(Extended Data fig. 5D). The variations within the HC group are less pronounced compared to those in the unhealthy group, thereby not impacting the results of unsupervised clustering(Fig. 2, D-G).

### 2.3 Speculation on the individual’s overall immune status

The model holds practical value only when the results can reflect an individual’s overall immune status. As this method is distribution based, so we expect the clustering result to be independent of sample size, provided that a certain threshold is met. Multiple sub-datasets with varying size were resampled from single sequencing data and the sample distance between them were calculated, we found the sub-datasets and the overall population holds the same distribution and the distance between them diminishes at a power-law rate with the increase of the sample size, a tenfold increase in data volume corresponds to a single-digit enhancement in the precision of estimating large samples from small samples(Fig. 3, A-C), indicating that a small data could represent the overall repertoire status of an individual. After calculating the average distance between CRC patients and HC group, we found the minimum amount of sample size to tell whether an individual is healthy or not at 10^4^ level. Clustering results with all samples’ data size reduced to 1∗10^4^ confirmed our computational results(Fig. 3D, Extended Data Fig. 7). Although the error is amplified due to the reduction in sample size, the unsupervised clustering of each health condition still shows obvious grouping. KD group and Infection group could still be distinguished. Naive HC, Post-IV HC and child group still holds unique clusters. These findings confirm the feasibility of the method to differentiate various immune statuses with minimal sample quantities. Data size normalization is not necessary for repertoire analysis in this method, datasets could compare to each other directly regardless of sample size as they all can represent the overall macroscopic status of the individual. Differences calculated from datasets with varying sample size ranging from 20k to 120k matched with the normalized situation and clustering results still aligned with clinical groupings(Fig. 2, B-G). This gives the feasibility to estimate the overall health status of an individual from small amounts of peripheral blood, even in the level of droplets.

**Fig. 3.**
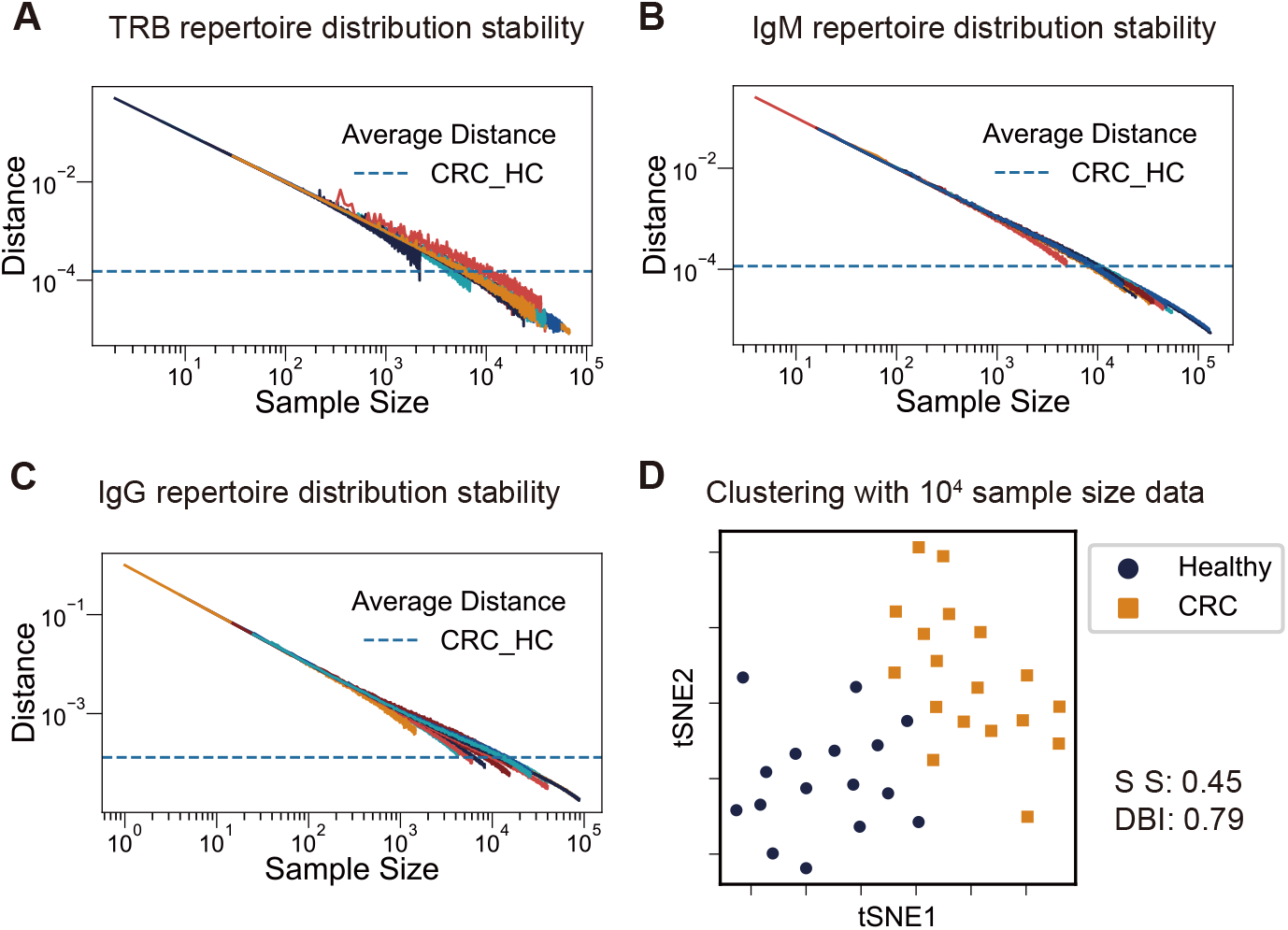
Structural characteristics of repertoire. Each line or point represents an individual. (**A, B, C**) The distances between the overall sequenced dataset and a small data subsets resampled from it are calculated. Distances decrease as the sample size increase. Dashed lines are the average distance between healthy adults and cancer patients. The discrepancy between the data subset and the overall population decreases at a power-law rate with the increase of the sample size in IgM, IgG and TRB. (**D**) data clustering of CRC and HC samples. data size are limited to 1∗10^4^, the clustering result are consistent with that of full sample size.

## 3 Discussion

In this article, we provide a generative markov method to demonstrate repertoire and quantify repertoire shift. By using a scale-free network model, we modeled the cell proliferation and SHM process into a preferential attachment model. This model gives a dynamic description of immune repertoire which is accurate enough to tell the difference between various health conditions as well as dynamic changes during immune process. Rather than depicting a static status of repertoire with indicators or summarize empirical rules from sequencing data and then verify their false positive rate, we focus on mathematical descriptions and provide comprehensive, quantitative tools. Importantly, this method could shift the paradigm from observing diseases to predicting specific illnesses based on immune characteristics. Immunological responses exhibit a cascade amplification effect, where minute quantities of an antigen can induce significant alterations in the immune repertoire. Compared to existence-based disease screening markers like cancer proteins and ctDNA, our distribution-based approach assesses alterations in immune distribution, thereby simplifying testing complexity and reducing the necessary sample volume. All this ensures its effectiveness across various health states of species and enables the representation of an individual’s overall health condition based on small sample sizes.

Traditional statistical methods can only compare the differences between the experimental group and the control group but are unable to explain why the measurements of the control group are considered normal. Our method provides an explanation for what constitutes ”normal” values by examining the process of formation, leading to a deeper understanding of the adaptive immune response.

Considering the statistics property of immune system, another model using the state of ideal gas can also get identical result with that from the scale-free model. Memory cells may be seen as gas molecules with smaller mass, as it have higher activation rate when giving same amount of stimulation. Many approximations were made in both mechanical and thermodynamic model to fit the biological process and made the model not so self-consistent, and the processes of B cell class switch, the generation of immune memory, and the calming process were not included in the model, which may cause inaccuracy in modeling the whole immune process. The model still showed significant accuracy in depicting and predicting the repertoire process in several diseases dynamically over time. So we believe the model shows accuracy at a certain degree and address such models based on phase transformation, grand canonical ensemble or dissipative process can further describe the immune process and will amendment the residual error.

We exclusively employed cost-effective bulk HTS sequencing, excluding the need for expensive single cell techniques in the analysis. Meanwhile, no clinical infomation are used in the study and the research results were obtained under blind conditions. These guaranteeing its extensive accessibility and adaptability in clinical conditions. No specific other method was coupled with the quantitative method including wetwork library preparation or sequencing method. Any BCR heavy chain or TCR *β* chain data containing read counts and mutation info(for BCR) are suitable for the method. The data in the study is derived from natural population cohorts, aiming to guarantee that the demographic profile within the sample mirrors the actual population distribution. Artificial filtering of the sample based on criteria such as age, sex or gender would diminish the universal applicability of the methodology. Therefore, these parameters were not used as criteria for sample selection.

In conclusion, we depicted the collective behavior of repertoire including B and T cells in activated immune response by modeling basic immunological cognition with mathematical distributions, and defined repertoire difference as the minimum efforts needed to transform a repertoire distribution to another. The immune system itself can be used as an indicator to detect fluctuations in changes in health status, the model may help solve the dilemma of early disease screening, misdiagnosis, disease stages identifications, potential secondary complications, and disease progression detection. The evidence-based evaluation of treatments and vaccines to regulate immune system in sick and healthy individuals can also be evaluated by monitoring the different repertoire pattern triggered. The consequent research will focus on the repertoire change in specific diseases, as the model’s availability to each kind of situation should be proved independently.

## 4 Methods

### 4.1 Materials

C57BL/6 mice were purchased from GemPharmatech(Nanjing, China), The VLP capsid protein(Escherichia virus Qbeta, a.a.1-133, Uniprot Id P03615) was purchased from Nanjing Liheng Biotechnology Co., Ltd.

### 4.2 CAWS Preparation

The CAWS were prepared from Candida albicans strain NBRC1385(Stock et al, 2016; Zhang et al, 2018; Tada et al, 2008), Candida albicans were incubated in C-limiting medium at 26*^◦^*C for 2 days at 160 rpm. Add equal volume of ethanol and overnight in 4*^◦^*C cold storage. Then collect the culture by centrifugation, dissolve the precipitate in water, stir for 2 hours, centrifuge the mixture again, harvest the soluble part and mix it with an equal volume of ethanol, and let it stand overnight. Finally, centrifuge the mixture to obtain the precipitate and dry it with acetone. Dissolve the CAWS dried product in 0.9% physiological saline and sterilize under high pressure for later use.

### 4.3 Immunization and sample collection

50ug VLP or 2mg CAWS are dissolved in PBS and given to each mouse through intraperitoneal injection. Mice are sacrificed at 3, 7, 14, and 28 days following VLP injection, and at 14 days following CAWS injection. Spleens were collected and homogenized to extract RNA. 3-5ml peripheral blood samples are collected from the hospital, and after isolating mononuclear cells, total RNA is extracted.

### 4.4 Wetwork Processing and HTS Sequencing

The wetwork and sequencing procedures(Mamedov et al, 2013) were performed by the GENEWIZ, lnc. Total RNA was extracted from the sample using Trizol(Takara, RNAiso Plus) according to the user manual. Template Switch PCR with UMI was performed for reverse transcription. The cDNA products were amplified by PCR with corresponding specific primers. PCR products were purified by DNA magnetic beads. Sequencing was carried out using an Illumina 2x250 pari-end configuration. IgA, IgM, IgG, TCR*β* chain data are sequenced independently

### 4.5 Data Preprocessing

Data preprocessing workflow includes QC, trim, alignment, annotation, and UMI deduplicating process. Raw seuqences are first checked by FastQC(Andrews et al, 2012), then trimmed by Trimmomatic(Bolger et al, 2014)(Q value ≥ 20). Alignments are performed by BLAST and annotated to the IMGT reference database(Lefranc et al, 2005). Sequences with distances less than 3bp as well as same UMI, V allele, J allele, and CDR3 length are considered as PCR duplicates. Only the mutations in the V regions are counted for accuracy.

### 4.6 Parameters in Preferential Attachment Model

Six parameters(four for TCR) are set to determine an unique immune repertoire, including (i) the distribution of initial stimulating signal from antigens in B and T cell differentiation from stem cells, (ii) the initial number of B or T cells involved in a particular immune response, (iii) the SHM rate in BCR which change stimulating signal randomly, (iv) the amplitude of change in stimulating signal caused by SHM, (v) the positive selection and the formulation to calculate the proliferation speed from stimulating signal. (vi) the overall increasion in the number of cells and clones through cell division and mutation.

How these parameters influences the repertoire are as follows:

(i) The distribution of initial stimulating signal should be a Gaussian distribution *N* (0, *σ*^2^) in the first quadrant, *σ* refer to the variance of initial stimulating signal. For the VDJ recombination process is highly randomized and should not have preference for any specific antigen(Extended Data Fig. 3A).
(ii) This paramater defines the number of cells at the beginning of simulation, which follow the distribution in (i). Small initiating cell number will generate a repertoire with concave rank-frequency distribution, larger initial cell number generate a repertoire with power-law rank-frequency distribution(Extended Data Fig. 3, B-C).
(iii) The SHM process is simulated by adding random changes in the stimulating signal, the non-lethal effective mutation rate in SHM is set to 0.7∗10*^−^*^3^ bp*^−^*^1^ division*^−^*^1^, and calculated to 0.24 cell*^−^*^1^ division *^−^*^1^ by Poisson distribution(Kleinstein et al, 2003). We set B cells with larger mutation numbers to have lower mutation rate and the maximum mutation number to 50(Allen et al, 1987). The occurrence of multiple mutations at the same base position are ignored in this model.
(iv) Mutation occurs in random positions on BCR gene, with no specific directions, so the change of stimulating signal should also follow a Gaussian distribution.
(v) Beacuse of the cellular apoptosis and selection process, high-affinity B and T cells are positively selected(Mayer et al, 2017), so in this model, the slope of the curve representing the relationship between the stimulating signal and proliferation speed must be greater than 1 in the positive x-axis. We tested the formulation with higher-order function, exponential function and composition function, all forms of functions can produce similar scale-free curve, the slope of the scale-free curve is directly proportional to the slope of the proliferation speed curve(Extended Data Fig. 3D). We use the exponential function *f* (*x*) = *e^x^*to generate the repertoire.
(vi) The cell proliferation and SHM process continues to occur when there is an immune stimulus, so a larger number of simulating cycles are repeated to simulate a more sustained immune response. As the number of cells increases, a power-law distribution emerges. As the number of cells increases, the slope of the curve also increases until it reaches a plateau(Extended Data Fig. 3, E-F).

The probability *P* (*i*) for a new cell to join a clone depends on the existing cell number *N* and proliferation speed *f* (*i*) of the clone, which is:

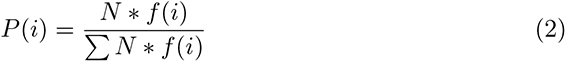

where Σ *N* ∗ *f* (*i*) is the partition function.

### 4.7 Ideal Gas Model

We assume the clonal expansion and somatic hypermutation process follows basic physics assumptions: The immune system maintains a maximum entropy distribution throughout the adaptive immune process(Mora et al, 2010); immune system evolve along the path of minimum energy consumption while maintaining maximum entropy. As different B or T cell clones proliferates with different speed and undergo different cycles of amplifications, Poisson distribution is not suitable to depict their collective behavior. so we then consider the mutation number *N*_Mut_ as a coarse-graining estimation of mutation velocity *v*, which follows the Gaussian distribution in each direction. Mutations are used as a metric parameter for clone shiftiness, and is a vector with at least two directional components, as the behavior of repertoire showing affinity maturation and epitope spreading process. B cells are considered as gas molecules and do not have interactions between similar cells.

We assume the maximum likelihood distribution between clone number and mutation number are the Maxwell-Boltzmann distribution(Mora and Walczak, 2016) and follow the state of ideal gas. So the projection of the 3D structure on the X-Y plane follows the ideal gas model(Extended Data Fig. 4). The energy of each molecule is 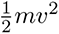, which is proportional to the square of mutation number, and total energy of the repertoire are the energy sum of each clone, which is identical with the result in Equation (1).

Neither early B cells nor T cells have got mutations, in order to expand the usability of the model to these cells, We set the ground energy state of each molecule to 0.1. The differences between two immune systems are calculated as the Wasserstein Distance between two repertoire distributions, which derives identical results with scale-free model.

## 5 Acknowledgments

## 5.2 Author contributions

Conceptualization, Data curation, Formal analysis, Software, Validation, Visualization, Writing - original draft: YC

Funding acquisition: PC

Investigation: YC, YL, JM, CH

Methodology: YC, SY, XX

Project administration and Supervision: PC, YC

Resources: PC, HN, HL, ML, YC

Writing - review & editing: PC, YC, XX

## 6 Ethical approval declarations

All mice were kept in SPF environment and fed ad lib in Jiangsu Province Academy of Traditional Chinese Medicine. All animal studies have been approved by Ethics Committee of Nanjing University of Chinese Medicine and performed in accordance with the ethical standards(AEWC-20220329-198).

Studies for the colorectal cancer and healthy adults have been approved by Ethics Committee of Nanjing Hospital of Traditional Chinese Medicine(KY2024025) Studies for the Kawasaki Disease patients and healthy children have been approved by Ethics Committee of Soochow University(SUDA20220906A01)

Methods were carried out in accordance with the relevant guidelines and regulations, Informed consent was obtained from all participants.

### 6.1 The inclusion and exclusion criteria

The samples in the study were selected from natural population cohorts, whth the criteria below:

1. Initial diagnosis of the corresponding disease for study inclusion;
2. Participants voluntarily join this study and sign an Informed Consent Form (ICF);
3. Exclude participants with a history of cancer;
4. Exclude participants whose immune function is affected by long-term medication use;
5. Exclude participants diagnosed with more than one primary disease simultaneously.
6. Exclude participants with pregnant or lactating women;
7. Exclude participants who have previously participated in other clinical trials and have not yet completed the trial;

## 7 Data and Code Availability

All code are avaliable at https://github.com/CYX-SAMIR/IR-simulation. All mouse data are avaliable at []. clinical data will be deposited in China National Genomics Data Center. All clinical data are avaliable at [].

---

**Extended Data Fig. 1.**
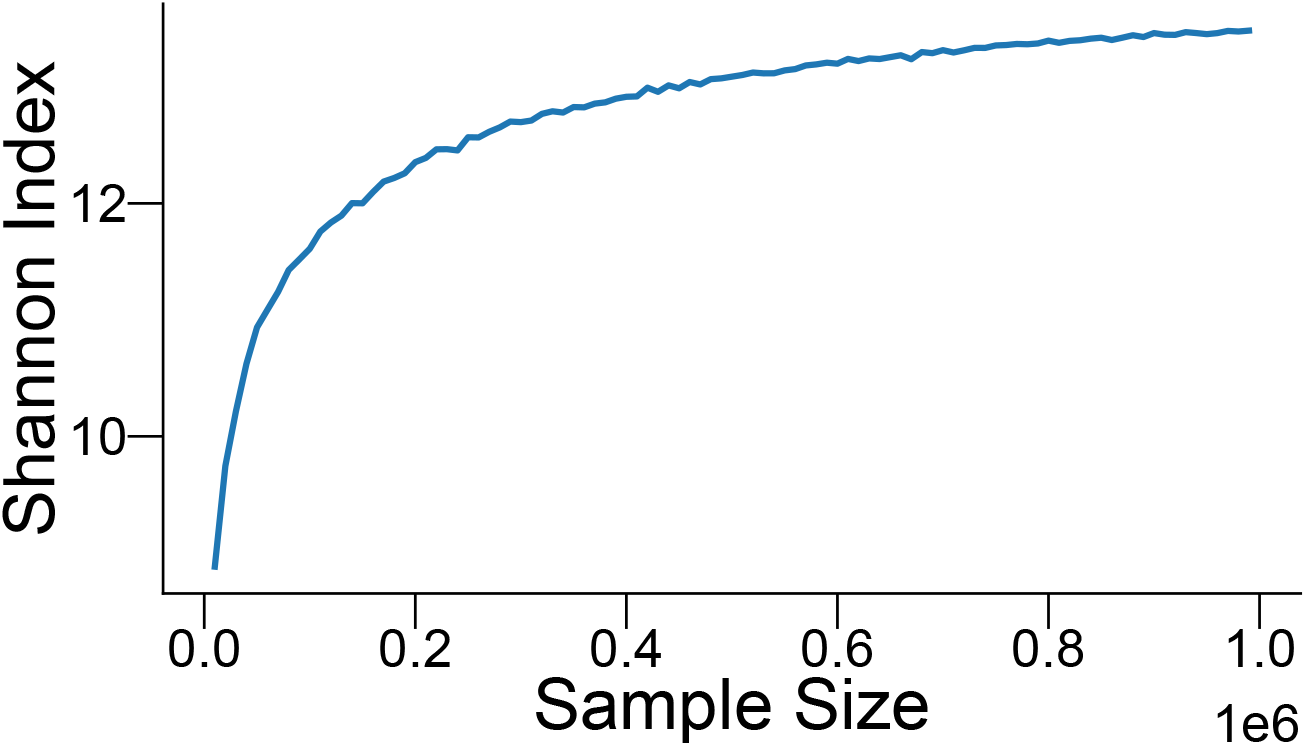
Correlation between Shannon Index and sample size. The Shannon Index value increases with the augmentation of the sample size.

**Extended Data Fig. 2.**
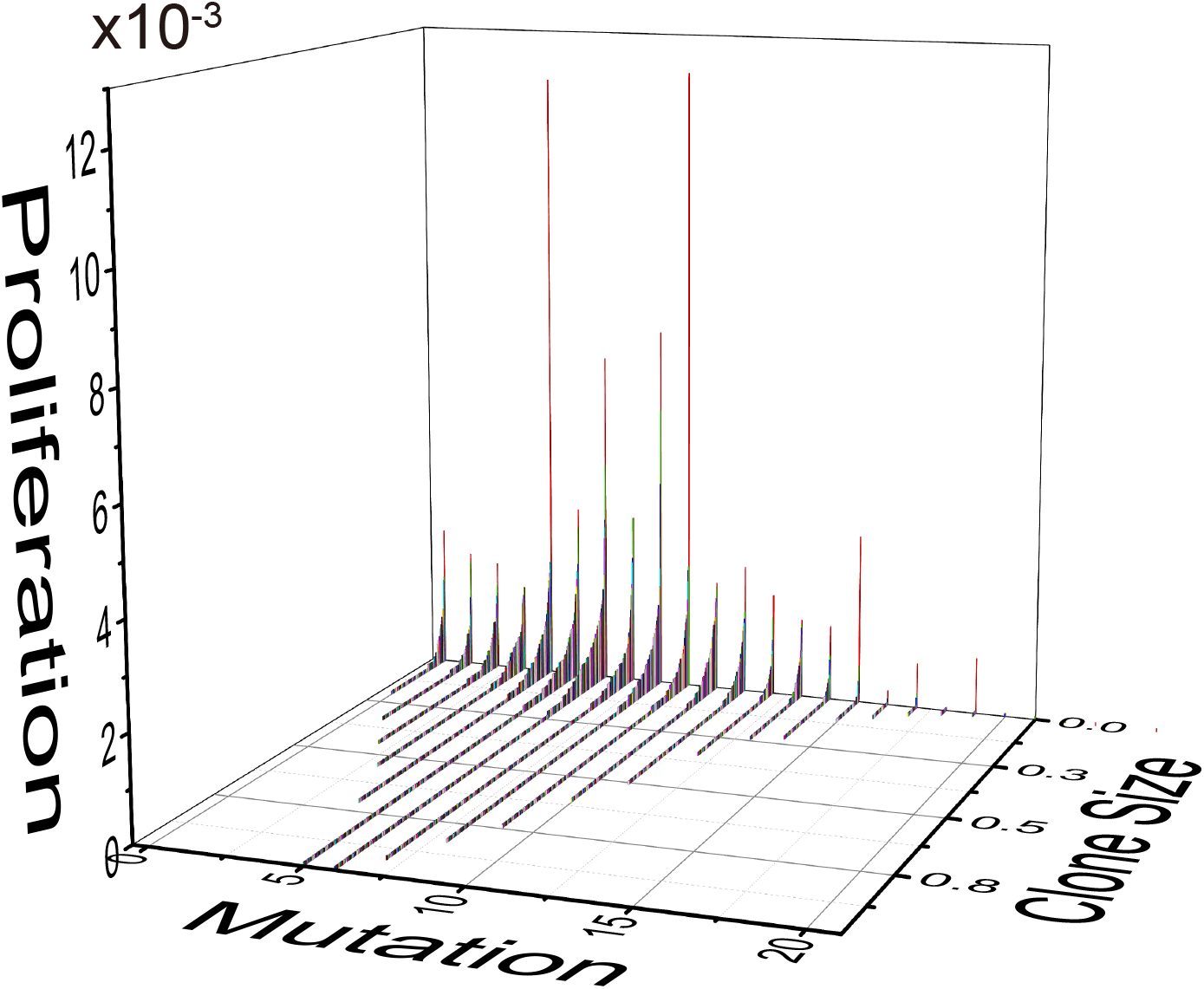
A 3D structural demonstration of repertoire. The three basis of the vector space are mutation, proliferation and clone number, each cell in repertoire occupies a point in the space based on its BCR or TCR sequence. The 3D map is uniquely determined by a repertoire.

**Extended Data Fig. 3.**
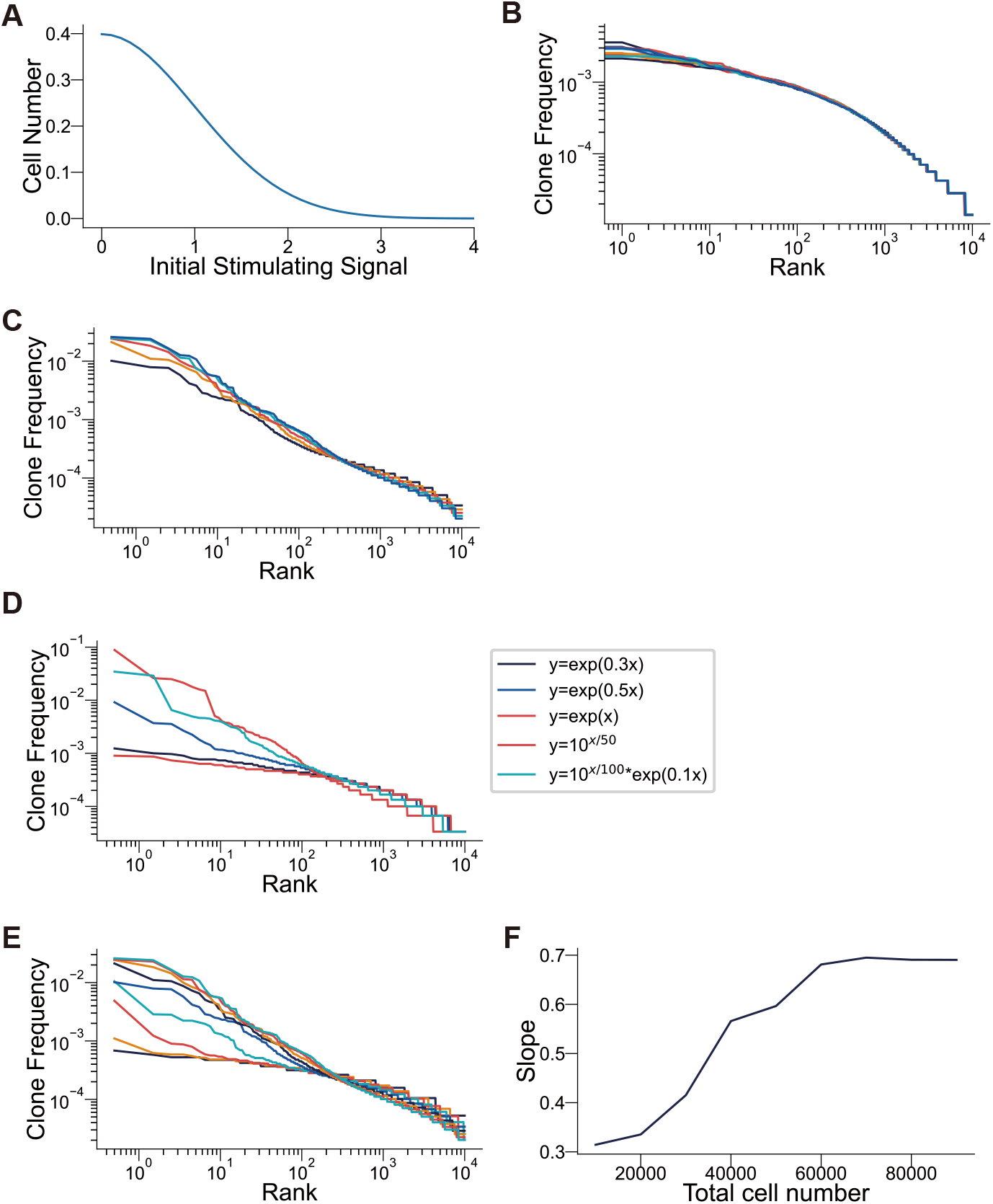
Parameters affecting the scale-free distribution of simulated repertoire. (**A**) The number of naive B or T cells with varying binding capacity and generate stimulating signal of different strength to a random antigen should follow a Gaussian distribution. (**B, C**) Simulation starts with 1k-5k cells and proliferate to 200k cells will generate a concave distribution, while power-law distribution emerges when starting with 50-90k cells. (**D**) Different formulations calculating the proliferation speed from stimulating signal can all produce a power law distribution, the greater the derivative of the function, the steeper the corresponding curve is. (**E, F**) The scale-free distribution generated from different final cell numbers range from 10k-100k. As the number of cells increases, the slope of the curve also increases until it reaches a plateau.

**Extended Data Fig. 4.**
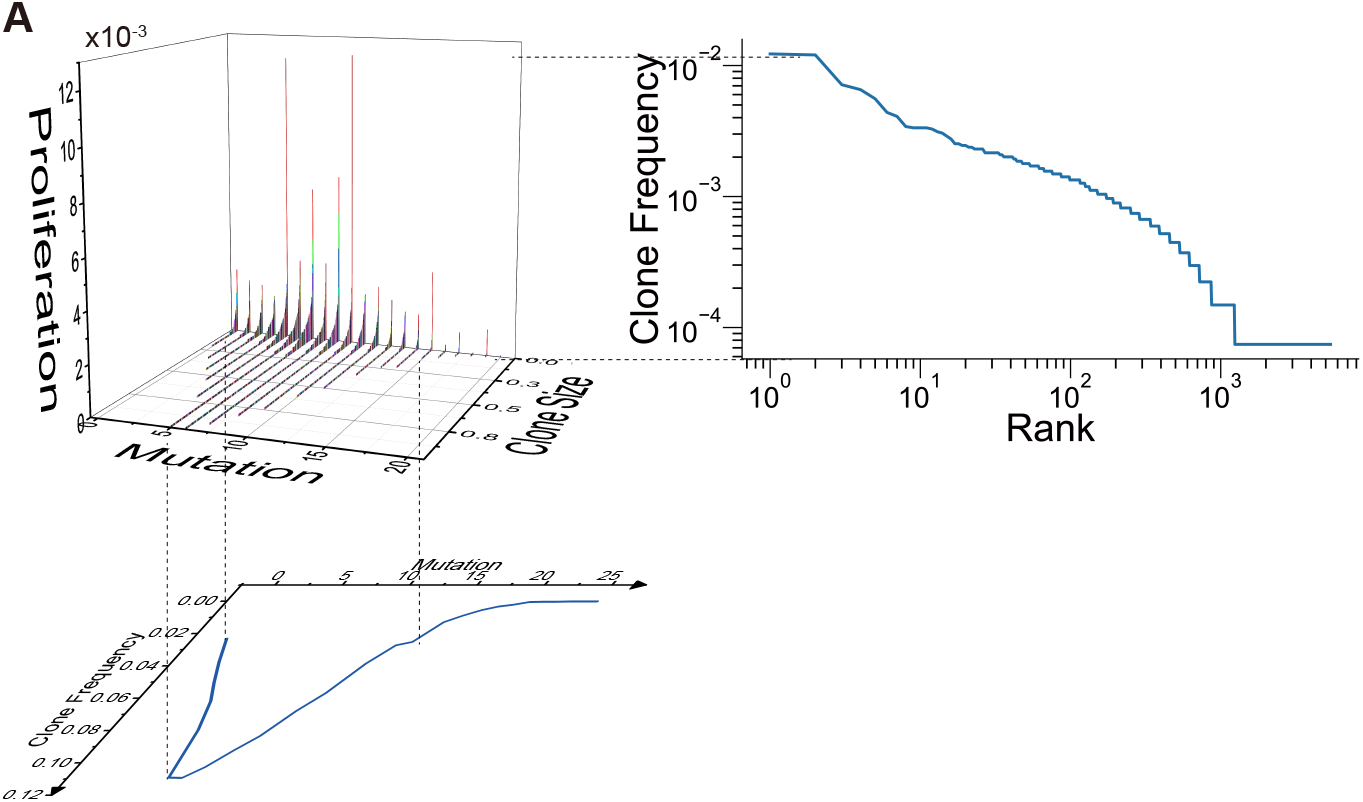
Correlation between 3D structures and various distributions. The dimensionality reduction results of the 3D structure at different angles exhibit the unimodal distribution and the scale-free distribution.

**Extended Data Fig. 5.**
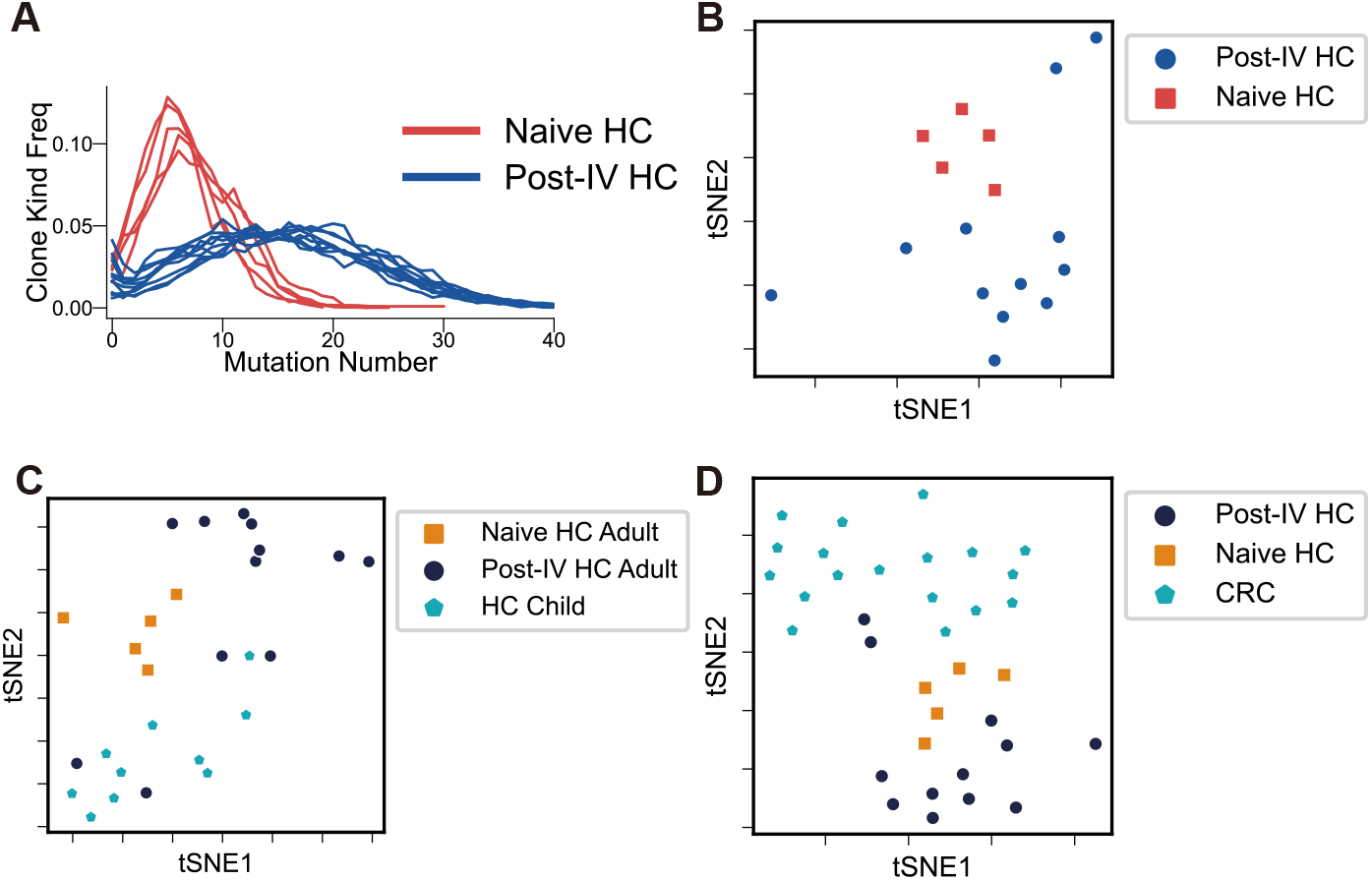
Subgrouping within the healthy group. (**A**) The mutation accumulation difference between naive healthy and Post-IV groups. (**B, C**) The unsupervised clustering of naive healthy, Post-IV group and healthy children group. Post-IV group exhibits fluctuations in clustering due to its diverse immune statuses. (**D**) In comparison to cancer patients, 14 out of 17 healthy individuals exhibited consistent immune statuses, while the remaining 3 displayed immune conditions that were on the boundary between the two states.

**Extended Data Fig. 6.**
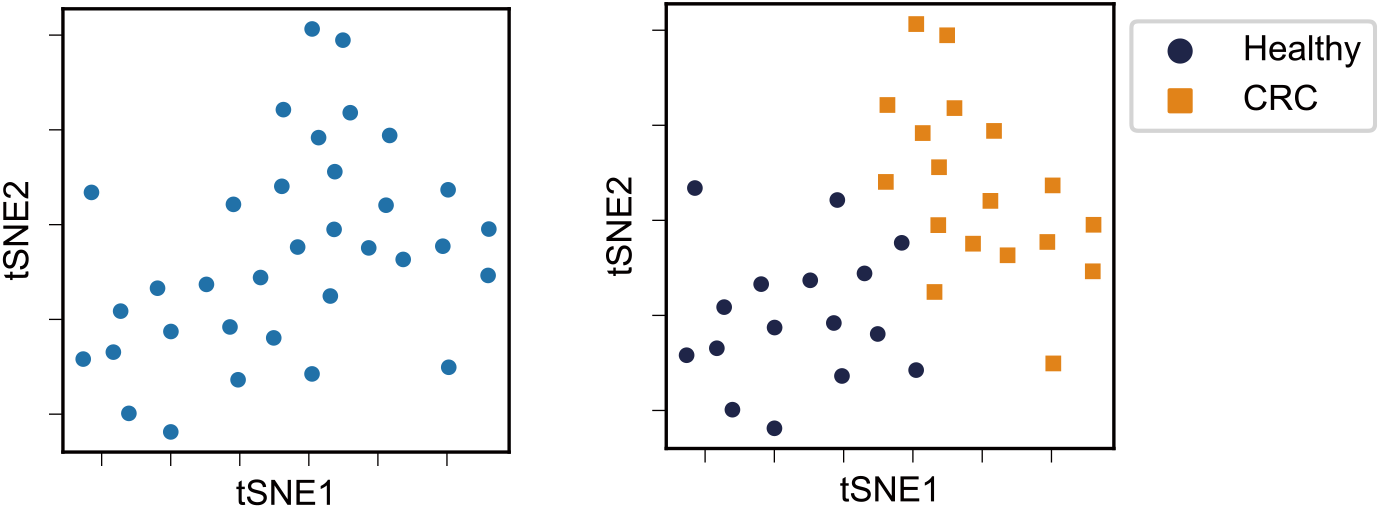
Blind clustering of repertoire data. The distances between any two samples were calculated and visualized to produce clustering results. The clinical information of HC and CRC are treated as a blind factor and are subsequently incorporated to verify the accuracy of the cluster.

**Extended Data Fig. 7.**
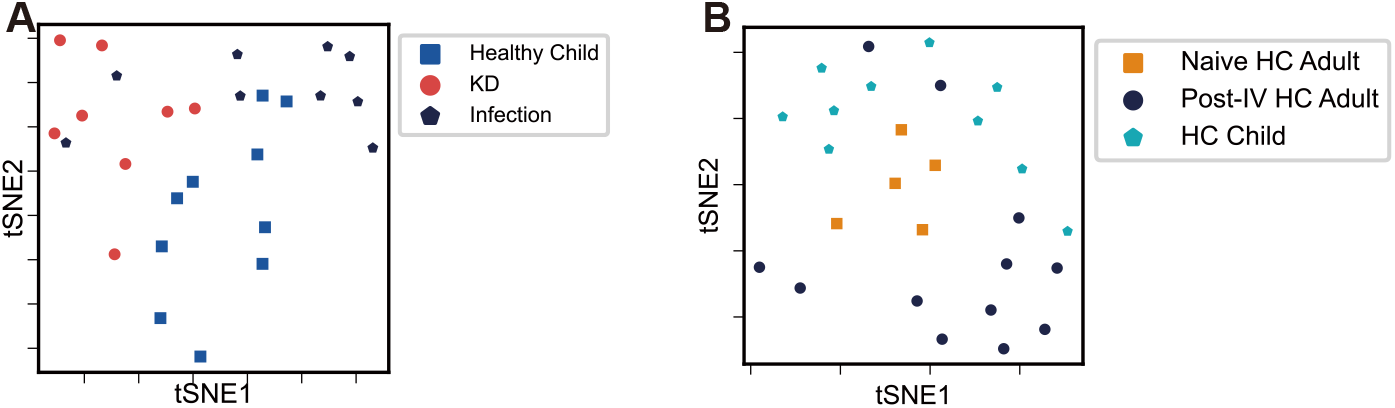
detailed clustering result of clinical data with sample size reduced to 1∗**10**^4^. (**A**) Distinct difference were observed between KD and Infection group with minimal sample size, the healthy child group demonstrates variations attributed to the persistent immune responses. (**B** difference among naive HC, Post-IV HC and HC child could also be observed with minimal sample size.)

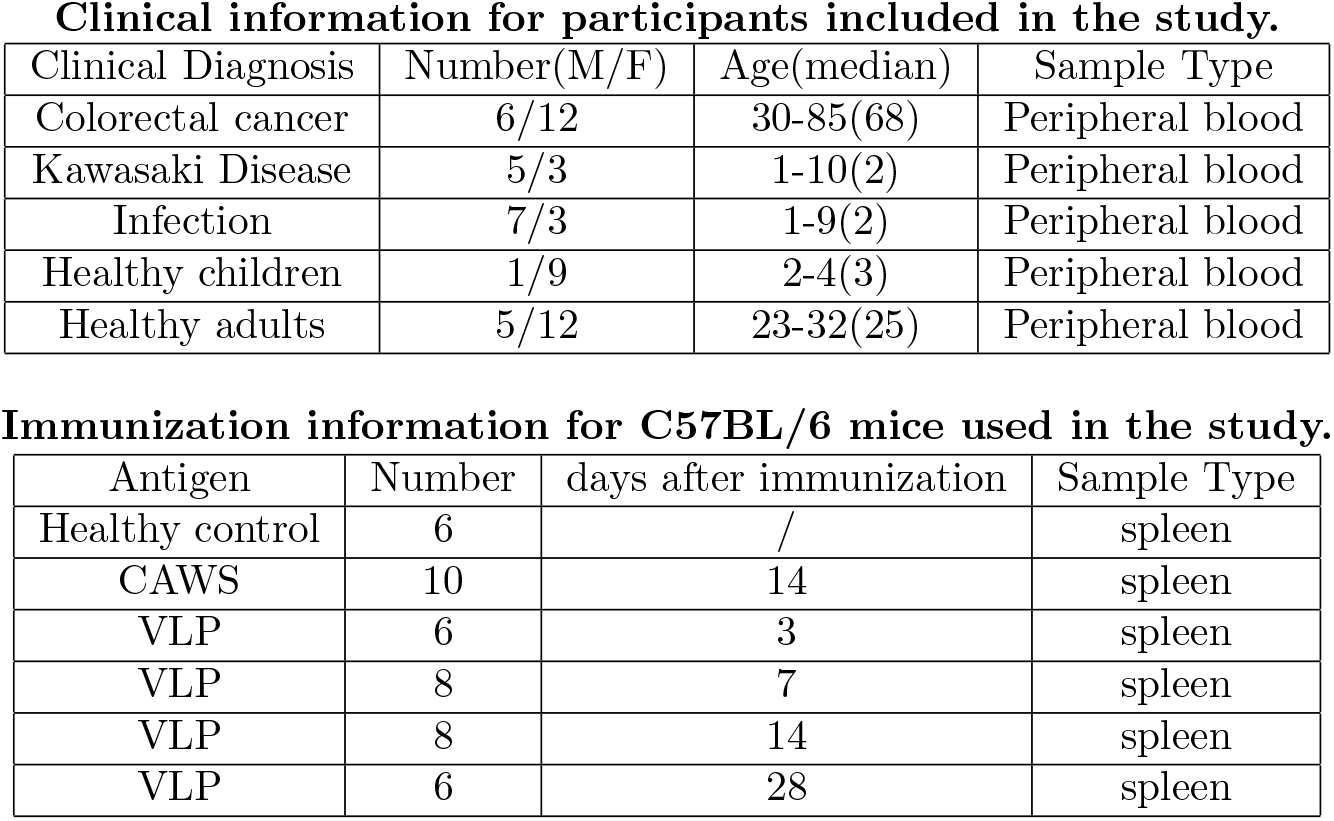

